# Exploring the limits of ComBat method for multi-site diffusion MRI harmonization

**DOI:** 10.1101/2020.11.20.390120

**Authors:** Suheyla Cetin-Karayumak, Katharina Stegmayer, Sebastian Walther, Philip R. Szeszko, Tim Crow, Anthony James, Matcheri Keshavan, Marek Kubicki, Yogesh Rathi

## Abstract

The findings from diffusion-weighted magnetic resonance imaging (dMRI) studies often show inconsistent and sometimes contradictory results due to small sample sizes as well as differences in acquisition parameters and pre-/post-processing methods. To address these challenges, collaborative multi-site initiatives have provided an opportunity to collect larger and more diverse groups of subjects, including those with neuropsychiatric disorders, leading to increased power and findings that may be more representative at the group and individual level. With the availability of these datasets openly, the ability of joint analysis of multi-site dMRI data has become more important than ever. However, intrinsic- or acquisition-related variability in scanner models, acquisition protocols, and reconstruction settings hinder pooling multi-site dMRI directly. One powerful and fast statistical harmonization method called ComBat (https://github.com/Jfortin1/ComBatHarmonization) was developed to mitigate the “batch effect” in gene expression microarray data and was adapted for multi-site dMRI harmonization to reduce scanner/site effect. Our goal is to evaluate this commonly used harmonization approach using a large diffusion MRI dataset involving 542 individuals from 5 sites. We investigated two important aspects of using ComBat for harmonization of fractional anisotropy (FA) across sites: First, we assessed how well ComBat preserves the inter-subject biological variability (measured by the effect sizes of between-group FA differences) after harmonization. Second, we evaluated the effect of minor differences in pre-processing on ComBat’s performance. While the majority of effect sizes are mostly preserved in some sites after harmonization, they are not well-preserved at other sites where non-linear scanner contributions exist. Further, even minor differences in pre-processing can yield unwanted effects during ComBat harmonization. Thus, our findings suggest paying careful attention to the data being harmonized as well as using the same processing pipeline while using ComBat for data harmonization.

## Introduction

Diffusion-weighted magnetic resonance imaging (dMRI) is the only non-invasive method that can map the living human brain’s structural connections. The measures derived from dMRI are quite sensitive to the subtle changes of the underlying microstructural tissue and have contributed to a wealth of knowledge about the abnormalities in several neurological and psychiatric disorders ^1,2^. However, results from neuroimaging studies often show inconsistent and sometimes contradictory results due to small sample sizes as well as differences in acquisition parameters and processing methods ^3,4^. Large collaborative efforts have led to Big-Data initiatives aimed at improving the poor reproducibility of small neuroimaging studies and gaining statistical power to detect subtle changes in neuropsychiatric disorders ^5–9^. However, scanner induced bias in dMRI signal limits direct pooling of the multi-site dMRI data.

*“Harmonization”* is a way to mitigate the measurement differences attributed to the scanner-, protocol-, or other site-related differences ^10–13^. Harmonization of multi-site dMRI datasets can dramatically increase the statistical power of neuroimaging studies and enable comparative studies pertaining to several brain disorders ^8,14^. When harmonizing dMRI data from different sites or scanners, one is ultimately interested in preserving the variability purely associated with the biology or disease, and, at the same time, removing other sources of variability (i.e., intrinsic or acquisition-related variability of the scanners), which can obscure the effect of interest.

Apart from the above-mentioned variabilities, the problem of harmonizing datasets from multiple sites becomes even more challenging when one considers the variability that can arise at each step of the processing pipeline. Typical dMRI pre-processing pipeline involves multiple steps, such as denoising, correction for subject motion, and geometrical distortion correction due to eddy currents and susceptibility artifacts. However, no consensus thus far exists on which processing steps to include, in which order, and which software to use for each step. Thus, minor changes in the pre-processing pipeline can bias the harmonization algorithm.

Various data harmonization methods have been proposed in the literature. These methods can be grouped into two main categories depending on the stage at which the harmonization is performed:

1. *Harmonization of the dMRI data at the signal level:* These approaches use the dMRI data directly and apply the harmonization at the signal level early in the pre-processing pipeline, avoiding (or protecting against) additional bias downstream in data processing as well as subsequent data modeling. Any model fitting, tractography, or connectivity analysis can be applied in later stages of the processing pipeline after harmonization ^12,15–17^. On the other hand, the approaches in this category require groups of healthy subjects from multiple sites for learning scanner differences.
2. *Statistical harmonization of diffusion maps:* These methods have been used in multiple fields, from genetics to functional MRI and recently in dMRI ^5,18–22^. They use the desired diffusion measures from multiple sites/scanners as input and apply statistical data pooling at the final stage of the processing pipeline to reduce the unwanted inter-site effects ^5,21,23,24^. The state of the art approach in this category is ComBat, a batch-effect correction tool used in genomics ^22^, and adapted to diffusion MRI harmonization^21^. ComBat is a powerful and fast statistical data pooling tool, which can estimate an additive and a multiplicative site-effect parameter at each voxel, thus accounting for voxel-wise scanner differences. Combat harmonization can be applied per voxel but also over entire regions of interest.

While many studies emphasized the benefits of the harmonization at the signal level, there are multiple advantages of using statistical harmonization approaches especially when a sample size that guarantees a statistically representative sample from each scanner is included in the studies. Furthermore, statistical harmonization (e.g., ComBat) is fast and easy to employ, and a more practical option when only diffusion measures of individual subjects from different sites are available, but not necessarily the dMRI data. For example, multi-center studies and consortia might only be able to share the diffusion maps (i.e. fractional anisotropy (FA) maps) of individual subjects; which makes statistical harmonization the only way to remove site effects. Thus, statistical harmonization approaches are more commonly used, but are more vulnerable to the pre-processing effects. It is, therefore, very important to explore these approaches thoroughly. In this study, we explore the commonly used approach, ComBat, on its ability to harmonize multi-site diffusion maps (measures). It is a well-known fact that a good harmonization algorithm should preserve the inter-subject biological variability from each site while removing only the scanner-specific effects and be minimally sensitive to changes in the pre-processing steps. To test whether ComBat has these characteristics, we used dMRI data from five sites, including 542 individuals. We investigated two important aspects of using ComBat harmonization, which have not been well-examined in prior studies. 1) How well does ComBat preserve effect sizes when used for harmonization, and 2) What are the effects of using different software packages to estimate the diffusion tensor on the harmonization performance of ComBat. Consequently, this study aims to answer these questions by harmonizing FA across five sites using ComBat.

## Methods

### Participants and diffusion MRI data acquisition

This study utilized dMRI data collected from five sites. Overall, these included a group of schizophrenia patients comprising 291 individuals (mean [SD] age, 28.57 [13.02] years; 198 [68%] male), as well as a healthy control group comprising 251 individuals (mean [SD] age, 26.77 [12.09] years; 157 [63%] male). Refer to Table 1 for demographics and acquisition parameters of each site. IRB/Sample previously reported in Cetin-Karayumak et al., 2019 ^8^.

**Table 1.** All dMRI data were collected as part of separate, individual studies. Basic demographics and dMRI acquisition information of each site are provided in the table.

| **Name of the study/location** | **Sample #** | **Age (mean+/-std)** | **Gender** | **dMRI data** |
| --- | --- | --- | --- | --- |
| **Site 1**<br>Siemens<br>Magnetom Trio 3T | 31H<br>46S | 40.74+/-11.31<br>38.37+/-11.43 | 22M, 9F<br>28M, 18F | b=1300 s/mm<sup>2</sup><br>42 directions<br>resolution=2x2x2 mm<sup>3</sup> |
| **Site 2**<br>GE Signa HDx | 70H<br>68S | 30.3+/-13.44<br>35.07+/-14.16 | 48M, 22F<br>51M, 17F | b=900 s/mm<sup>2</sup><br>59 directions<br>resolution=1.7x1.7x1.7 mm<sup>3</sup> |
| **Site 3**<br>GE HDx 3T | 69H<br>63S | 21.50+/-5.048<br>21.31+/-4.94 | 45M, 24F<br>46M, 17F | b=1000 s/mm<sup>2</sup><br>31 directions<br>resolution=0.94x0.94x2.5 mm<sup>3</sup> |
| **Site 4**<br>Siemens Tim Trio 3T | 25H<br>42S | 37.04+/-11.14<br>37.67+/-10.83 | 16M, 9F<br>32M, 10F | b=1000 s/mm<sup>2</sup><br>60 directions<br>resolution = 2x2x2 mm<sup>3</sup> |
| **Site 5**<br>Siemens Sonata 1.5T | 56H<br>72S | 16.5+/-1.74<br>17.17+/-1.66 | 26M, 30F<br>41M, 31F | b=1000 s/mm<sup>2</sup><br>60 directions<br>resolution=2.5x2.5x2.5 mm<sup>3</sup> |
| **Total** | 251H<br>291S | 26.77+/-12.09<br>28.57+/-13.02 | 157M,<br>94F<br>198M,<br>93F |  |

Abbreviations: H: Healthy Controls, S: Schizophrenia Patients; F: females; M: males.

### Diffusion MRI data pre-processing

A standardized processing procedure (eddy current and motion correction) was applied to all subjects using FSL’s eddy correction software ^25^. Next, for each dMRI data in 5 sites, we fit the diffusion tensor with ordinary least squares fitting and computed FA using three software packages (FSL v. 6.0.1 ^26^, MRtrix v. 3.0 ^27^, and Slicer v. 4.10.2 ^28^). To compare the whole brain white matter and major bundles across sites, the Illinois Institute of Technology (IIT) v. 4.1 Human Brain FA atlas in MNI space ^29^ was registered to the FA maps in subject space using *antsRegistrationSyn.sh ^30^*. Next, we warped the IIT white matter probabilistic atlas consisting of 17 bundles to the subject space. The probabilistic bundles were thresholded at 0.25, resulting in a total of 17 regions of interest (ROIs) of white matter fiber bundles. For each ROI, mean FA was computed by averaging FA across all voxels traversing the individual ROI. Additionally, average FA in the whole brain white matter was computed to complement the analysis. Mean FA of 17 ROIs + whole brain was used as input to ComBat to perform harmonization across the five sites.

### ComBat Harmonization

The ComBat model was introduced in the context of gene expression analysis by Johnson et al. ^22^ and reformulated by Fortin et al. ^21^ in the context of dMRI. Briefly, ComBat uses a Bayesian framework to estimate the additive and multiplicative site-effect hyperparameters at each voxel or ROI empirically. In this work, we utilized the Matlab version of ComBat software (https://github.com/Jfortin1/ComBatHarmonization) for harmonization of 17 ROIs + whole brain across five sites. Age, sex and disease status were added as biological covariates to the design matrix.

### Post-analysis

The ability of ComBat to perform accurate harmonization was tested by comparing the effect sizes between groups before and after harmonization. Effect sizes for FA differences in Cohen’s d were computed between healthy females and males, as well as separately between all patients and controls to assess group differences. The mathematical definition of effect sizes in Cohen’s d can be found in ^12,31,32^. According to Cohen’s d criteria, small effect size is 0.2, medium is 0.5 and large is 0.8. The effect sizes were computed before and after ComBat harmonization at each site for comparative analysis.

## Results

Inter-group/biological variability was measured by the effect sizes of the FA differences between two groups within each site:

(group i) healthy female and male controls;
(group ii) all healthy controls and schizophrenia patients.

We conducted the following two experiments to assess whether the inter-group variability was preserved after harmonization. We note that the effect sizes before and after harmonization are respectively referred to as “before” and “after” in figures.

### Experiment 1

In the first experiment, the entire dMRI data pre-processing was conducted in a consistent manner with the same software across five sites. We assessed the effect sizes of FA differences of females and males (group i) after harmonization, where FA was computed using FSL for all sites and mean FA of 17 ROIs + the whole brain was given as input to the ComBat harmonization algorithm. Figure 2 depicts the effect sizes of each ROI and whole brain for site 1 and site 5, both before harmonization and after harmonization with ComBat (refer to supplementary Figure 1 for the other sites). At site 1, the effect sizes were preserved before and after harmonization for nearly all ROIs, barring a reduction in effect size in the negative direction for left inferior fronto occipital fasciculus (before harmonization: −0.25, after harmonization: −0.05). On the other hand, for site 5, the effect sizes for almost all ROIs were altered after harmonization, i.e. the effect sizes were amplified across many ROIs after harmonization with ComBat (e.g. see right cingulum - cingulate gyrus portion, before harmonization: −0.2, after harmonization: −0.5). For five ROIs (right inferior fronto occipital fasciculus, right corticospinal tract, left superior longitudinal fasciculus, left corticospinal tract, left cingulum - hippocampal portion), the sign of the effect size was flipped, indicating that after harmonization the direction of the FA differences in group i were reversed.

**Figure 1.**
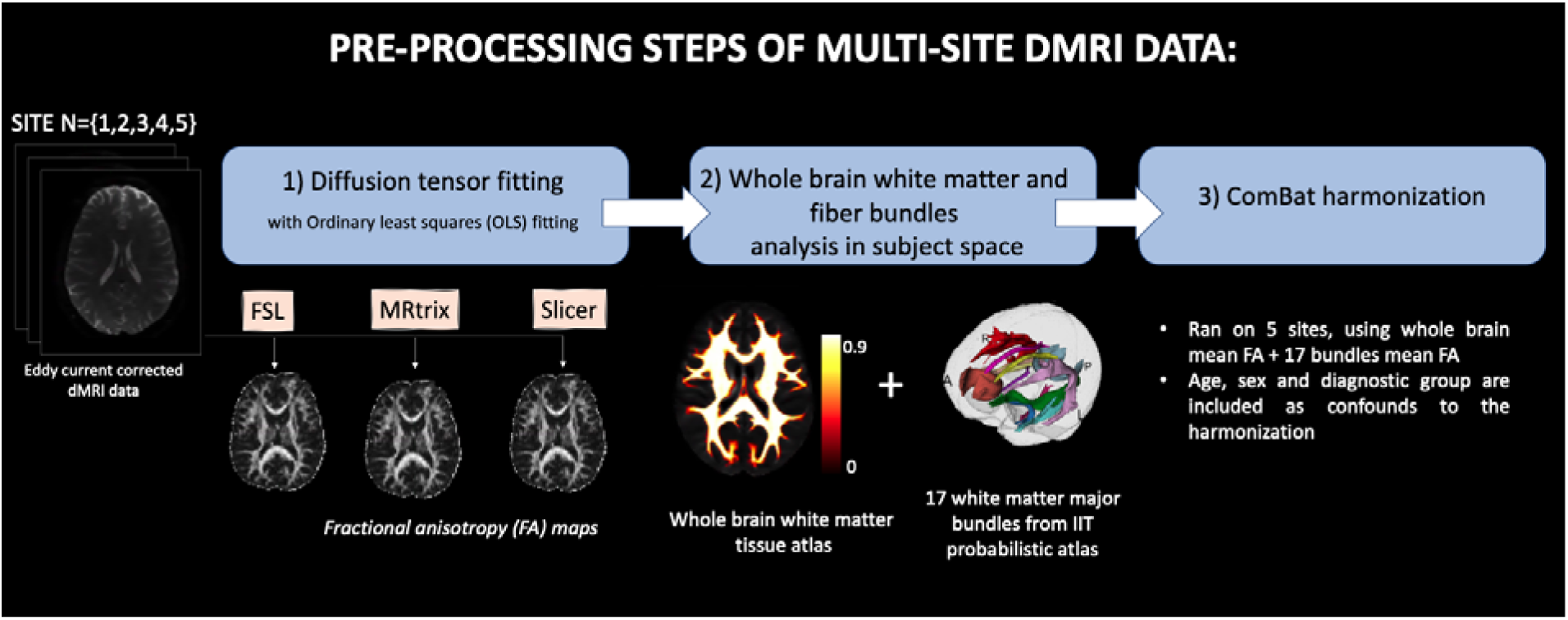
Pre-processing steps of dMRI data collected from 5 sites: (0) All dMRI data is axis aligned, centered and eddy current corrected using the same pipeline; (1) for all subjects, FA is computed using three software with ordinary least squares fitting; (2) FA atlas is registered to each subject space using antsRegistration and probabilistic maps for 17 major bundles and whole brain white matter tissue map is warped to the subject space; (3) mean FA of whole brain white matter and 17 bundles are computed and given as input to ComBat harmonization.

**Figure 2.**
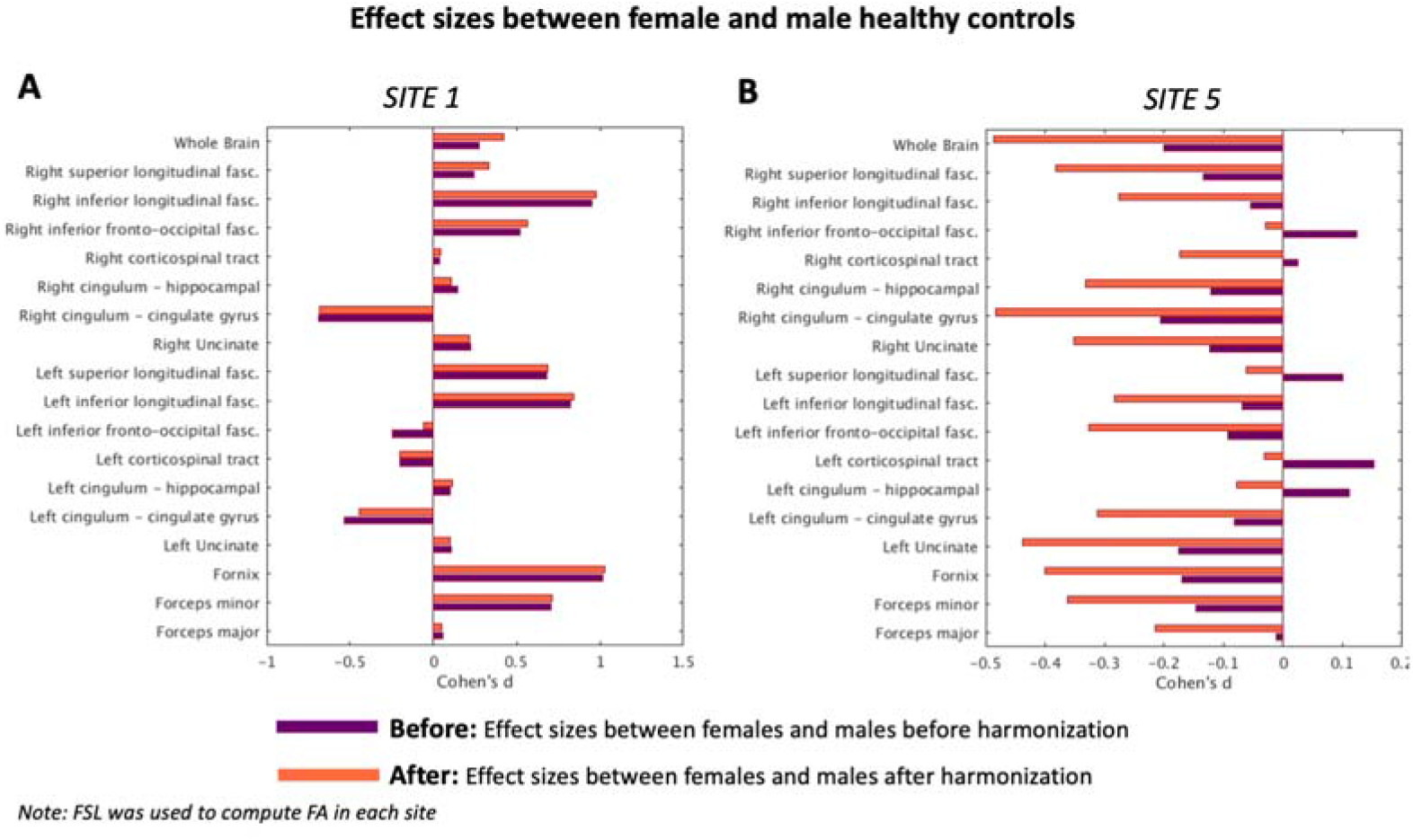
Effect sizes (Cohen’s d) of FA differences between female and male healthy controls are computed to assess whether the inter-subject variability is preserved after harmonization. The results are shown for site 1 and site 5 for the whole brain and 17 bundles. Purple bar represents effect sizes between two groups before harmonization (“before”). Orange bar represents effect sizes after harmonization, where FSL’s FA was given as input to the harmonization (“after”). Data pre-processing was done in a consistent manner with the same software. At site 1, the effect sizes were almost preserved before and after harmonization (barring a reduction in effect size for left-inferior-fronto-occipital-fasciculus). However, at site 5, we observe that the effect sizes for almost all fiber bundles are altered after harmonization. Notably, ComBat seems to have increased the effect sizes across many fiber bundles. Most importantly, in a few fiber bundles, the sign of the effect size is flipped, indicating that after harmonization the direction of the differences was reversed, which can dramatically reduce reliability of the results.

### Experiment 2

In the second experiment, we used different software packages to compute FA for each site to explore how subtle changes in the processing pipeline affect harmonization. The mean FA of 17 ROIs + the whole brain computed using different software packages for each site was provided as input to the ComBat harmonization algorithm. Figure 3 depicts the effect sizes between all healthy controls and schizophrenia patients (group ii) of site 3, both before harmonization and after harmonization with ComBat (refer to Supplementary Figure 2 for the other sites). We observed that when different software was used for each site to compute FA, effect sizes were not preserved after harmonization and the absolute differences in effect sizes between before and after harmonization were as large as 0.7. In addition, the effect sizes of seven ROIs changed sign (see right superior longitudinal fasciculus, right inferior longitudinal fasciculus, right cingulum hippocampal, right cingulum - cingulate gyrus, right uncinate, left cingulum - hippocampal, fornix). For example, for the right cingulum - hippocampal portion, while the effect size before harmonization was 0.3, it became −0.2 after harmonization.

**Figure 3.**
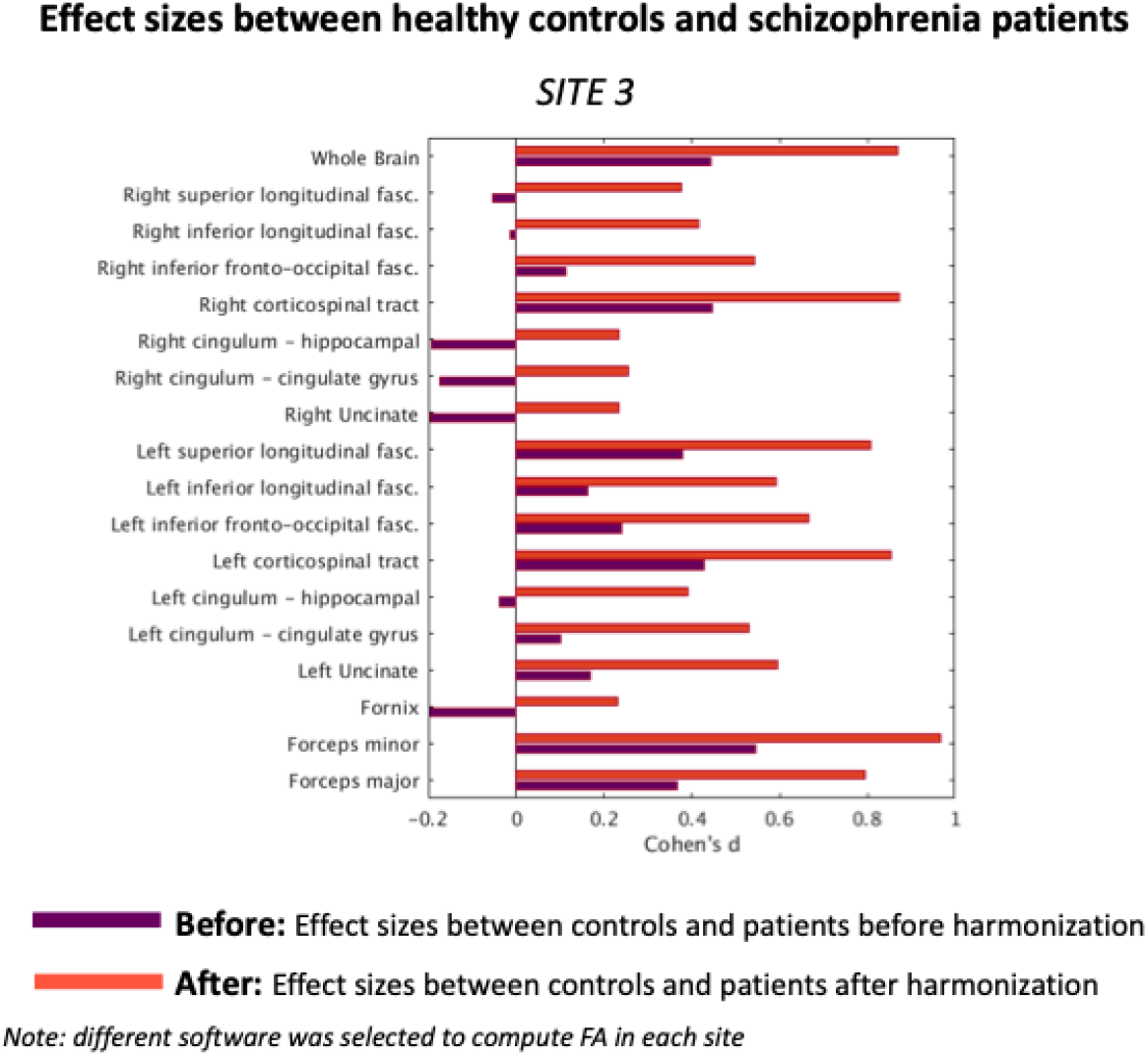
Effect sizes (Cohen’s d) of FA differences between healthy controls and schizophrenia patients are computed to assess whether the inter-subject variability is preserved after harmonization. The results are shown for Site 3 for the whole brain and 17 bundles. Purple bar represents effect sizes between two groups before harmonization (“before”). Orange bar represents effect sizes after harmonization, where FSL’s FA was given as input to the harmonization (“after”). To explore how subtle change in the pre-processing pipeline affects harmonization, different software to compute FA were given as input to the ComBat harmonization algorithm. After harmonization, effect sizes were not preserved such that absolute differences between “before” and “after” effect sizes were up to 0.7. In addition, we point out to the bundles where the effect sizes changed direction (e.g. right-cingulum-hippocampal). These results demonstrate that inter-subject variability was not preserved after harmonization and subtle changes in the pre-processing pipeline induced much larger biases, which were not properly accounted for by ComBat.

## Discussion

Harmonization of dMRI data along with a consistent processing pipeline is critical for reproducible neuroscientific research and provides increased statistical power. Statistical harmonization methods have been widely preferred in the literature for joint analysis of dMRI derived scalar maps due to their simplicity in neuroimaging applications. These harmonization methods such as Meta-analysis ^5^ and ComBat ^21^ statistically pool dMRI measures (FA, mean diffusivity, etc.) at the final stages of the processing pipeline. In this study, using a large dataset of 542 subjects from 5 sites, we evaluated ComBat in its ability to harmonize multi-site dMRI data by comparing the effect sizes (Cohen’s d) in two groups, before and after harmonization. To this end, we conducted two experiments. In both experiments, we applied the same pre-processing pipeline for all sites, except that in experiment 2, we used different software packages to fit the diffusion tensor and compute FA for each site.

In experiment 1, we observed that even when all processing steps, as well as diffusion tensor estimation software were identical for all sites, ComBat harmonization still seemed to alter the inter-group differences and, in some cases, flipped the direction of effect sizes. This indicates that after harmonization, the direction of the differences was reversed. This might be due to ComBat’s optimization procedure that assumes that the site-effect parameters follow a particular parametric prior distribution (Gaussian and Inverse-gamma), which might not be true in all scenarios. We note that our findings support results in structural neuroimaging studies demonstrating that the standard ComBat approach might not necessarily preserve biological variability ^33–35^. Additionally, we notice that at site 5, the data was acquired on a 1.5T scanner, whereas at site 1 it was acquired on a 3T scanner. Thus, the scanner differences are potentially highly non-linear which are not well captured by ComBat.

In experiment 2, we investigated the effects of additional sources of variability that can possibly arise at any step of the dMRI processing pipeline, when different software packages or versions are used for data pooling using ComBat (for example, when already processed data is shared for joint analysis across consortia or sites). We first applied FSL’s eddy to all sites for motion and eddy correction. We later used three different software packages (FSL, MRtrix, Slicer) to estimate the diffusion tensor and compute FA at different sites. We consistently used ordinary least squares fitting for diffusion tensor computation in all three software packages. These software packages perform pre-processing in slightly different ways and order (e.g., signal normalization by b0, denoising, thresholding after normalization to restrict data between 0 and 1, fixing negative eigenvalues, etc.). In this experiment, we observed that group differences were not preserved after harmonization. Thus, minor differences in pre-processing produced medium to large differences in the data after harmonization. This might be due to the fact that, the data at site 3 was interpolated on the scanner (in-plane resolution was 0.94 x 0.94 mm^2^), potentially causing the data to have many more negative values or negative eigenvalues and more partial volume effects due to smoothing. Consequently, minor differences in the pre-processing steps resulted in non-linear changes in the input data, thereby causing ComBat to perform in an unexpected manner.

We note that, we only evaluated the standard ComBat method^21^ that is available as open-source (https://github.com/Jfortin1/ComBatHarmonization). The recent versions of ComBat, such as the generalized additive model/GAM - ComBat ^34^ might mitigate some of the problems; however, it remains to be tested on large dMRI datasets like the one used in this work.

## Conclusion

This study thoroughly explored the powerful ComBat harmonization in a large dMRI data of 542 individuals. We demonstrated that while ComBat harmonization preserves the effect sizes of between-group FA differences in some sites, it alters the between-group differences (significantly in some cases) and in some ROIs flips the sign of the effect sizes. We thus suggest caution while using harmonization methods like ComBat. Most importantly, we recommend using a consistent pre-processing pipeline in multi-site dMRI studies, more specifically, not to combine dMRI measures processed with different software packages.

## Acknowledgments

The authors have been supported by the following grants: NIH R01 MH119222 (Rathi), R01 MH102377 and K24 MH110807 (Kubicki), Swiss National Science Foundation grant 152619 (Walther). The project also acknowledges that the research has partly been supported by the BWH Program for Interdisciplinary Neuroscience, through a gift from Lawrence and Tiina Rand (Cetin-Karayumak).

